# Genes acquired by horizontal gene transfer share common regulatory patterns in *Photorhabdus laumondii*

**DOI:** 10.1101/2023.10.31.564978

**Authors:** Lydia Rili, Marcus Simoes, Maythem Ali, Brandon L. Findlay

**Affiliations:** Department of Chemistry and Biochemistry, Concordia University, Montreal, Québec, Canada; Department of Biology, Concordia University, Montreal, Québec, Canada

**Keywords:** Natural product biosynthesis, horizontal gene transfer, alternative sigma factors, enhancer binding proteins, gene induction, *Photorhabdus laumondii* TTO1

## Abstract

Biosynthetic gene clusters are readily transferred between microbes through horizontal gene transfer, but to provide benefit to their host they must be induced to high levels during times of stress. Working with the insect pathogen *Photorhabdus laumondii*, here we study the effect of plasmid-induced stress on natural product biosynthesis. We find that the expression plasmid pARO190 alone was sufficient to induce production of an array of natural products, and that the induction could be increased with heterologous expression of either bacterial enhancer binding proteins or a random DNA sequence. Overall, this suggests that generalized stress may be an effective way to increase production of natural products in *P. laumondii*.

## Introduction

Driven by advances in genomic sequencing the field of natural product discovery is in a renaissance. Genomic sequencing has shown that even well studied *Streptomyces* strains express less than a quarter of their biosynthetic potential under laboratory conditions (Tanaka *et al*., 2013). The remaining biosynthetic gene clusters (BGC) are tightly repressed, and the vast majority of compounds they produce are uncharacterized. While industry largely ended their bioprospecting efforts in the 1990’s due to the repeated re-isolation of known bacteria and well-characterized natural products (Katz and Baltz, 2016), it appears that the tens of thousands of strains isolated during the golden age of antibiotics are far from exhausted. Improving our understanding of bacterial gene regulation may lead to the discovery of hundreds of thousands of new natural products.

Activating these silent BGC can be a significant undertaking. In many cases the key regulators that control these clusters are unknown, requiring either resource-intensive heterologous expression or combinatorial analysis of growth conditions and possible chemical inducers (the OSMAC approach) (Romano *et al*., 2018). However, it may be possible to sidestep these issues by taking advantage of the mobile nature of BGC themselves. BGC can be readily transferred between related and unrelated strains via horizontal gene transfer (HGT) (Jensen, 2016), a process that is widespread in terrestrial and marine environments (McDaniel *et al*., 2010; Aminov, 2011; Ziemert *et al*., 2014). We hypothesize that to maximize fitness these newly acquired BGC will be transcribed only when they provide the greatest advantage to their new host. Absent any knowledge of their function, this would require either evolutionary adaptation of the incoming regulatory sequences to match host paradigms, or non-specific activation of all acquired genes. The latter would require conserved regulatory sequences across nearly all BGC, as well as host transcription factors that recognize this skeleton key.

One such potential regulator was identified following transfer of the BGC for oxytetracycline from *Streptomyces rimosus* ATCC 10970 to the model bacterium *Escherichia coli* BAP1 (Stevens *et al*., 2013). Overexpression of a single protein, the sigma factor RpoN (σ^54^), was sufficient to activate the cluster and produce oxytetracycline in *E. coli*. This alternative sigma factor binds to RNA polymerase, determining its promoter specificity (Bush and Dixon, 2012). The consensus sequence for σ54 is well known, and a screen of BGCs in Actinibacteria, Firmicutes and Proteobacteria revealed that ∼70% of polyketide and non-ribosomal peptide synthase-containing clusters contained σ^54^ promoters, despite the passage of billions of years since the last common ancestor between these three *phyla* (Stevens *et al*., 2013).

However, in Proteobacteria σ^54^ is necessary but not sufficient for expression of genes from σ54 promoters. One of several bacterial enhancer binding proteins (bEBPs) are required to hydrolyze ATP and initiate transcription (Bush and Dixon, 2012). These bEBPs appear to direct activation of σ^54^-linked genes in response to stress, with roles in phage shock, nitrogen limitation, tyrosine biosynthesis, among others (Bush and Dixon, 2012). Most bEBPs consist of a three interlinked domain, with an N-terminal regulatory domain that activates the protein in response to a cognate kinase or chemical signal, a AAA+ ATPase domain for the initiation of transcription, and a C-terminal DNA binding domain that directs the bEBP to its intended genes. When expressed in isolation the central domain pan-activates σ^54^, and the core domain of the bEBP DctD from *Sinorhizobium meliloti*, tDctD, has been used to determine the σ^54^ regulon in *S. meliloti* and *Salmonella enterica* (Xu *et al*., 2004; Samuels *et al*., 2013).

Interested in the effect of bEBP overexpression on natural product biosynthesis, we set out to express host regulators in the well-characterized entonomopathogen *Photorhabdus laumondii* TTO1. Metabolite production was assessed via LC-MS, revealing activation of a number of BGCs that are poorly expressed during normal lab growth. Surprisingly, we also found that conjugation of a plasmid lacking bEBPs was sufficient to induce natural product biosynthesis, and that one plasmid with a malformed insert induced biosynthesis at levels on par with the most effective of the bEBPs.

## Materials and Methods

### Molecular biology

The strains and plasmids used in this study are listed in Table 1. *E. coli* MFDpir was a generous gift from Luis Ángel Fernández Herrero (Centro Nacional de Biotecnología, Madrid, Spain) (Ferrières *et al*., 2010), while *P. laumondii* TTO1 was purchased from DSMZ (Braunschweig, Germany). pARO190 was acquired from Cedarlane (Burlington, Canada). The coding sequence for the *Sinorhizobium meliloti* gene *dctD*_141-394_, with flanking KpnI and HindIII recognition sequences, was custom-synthesized as a single sequence by Gen9 alongside(see supplementary materials). Genes for each other bEBP were amplified from the *P. laumondii* TTO1 genome (see Table S1 for a list of primers), digested with KpnI and HindIII, then ligated into pARO190 with T4 DNA ligase. As *glrR* contained a KpnI site for this gene HindIII and SaII were used instead. Each plasmid was then transformed into *E. coli* MFDpir cells previously made competent with TSS buffer (Chung and Miller, 1993). The plasmids were then transferred into *Photorhabdus laumondii* TTO1 via conjugation. In brief, cells from both strains were grown overnight in TSB at 30 °C and 225 rpm (with *E. coli* MFDpir cells supplemented with 0.3 mM diaminopimelic acid (DAP) and 100 µg/L of carbenicillin). Cells were then diluted 1:100 into fresh media and grown until an OD_600_ of 0.4. Cells were centrifuged for 1 min at 13 krpm, the supernatant was discarded, and the cells were resuspended in fresh media at their former concentration. *E. coli* and *P. laumondii* were then mixed in a 1:3 ratio, and plated onto TSA plates supplemented with diaminopimelic acid. Following overnight incubation at 30 °C colonies were resuspended in TSB and plated directly onto TSA plates supplemented with 100 µg/L of carbenicillin but not DAP for a period of 48-72h at 30 °C. Repeated attempts to transform *P. laumondii* by chemical or electrical means were unsuccessful.

**Table 1.** Strains and plasmids used in this study.

| Strain | Notes |
| --- | --- |
| *Photorhabdus luminescens* TTO1 (DSM15139) | Wild type, formerly known as *Photorhabdus luminescens* subsp. *laumondii* TTO1. |
| *E. coli* MFD(*pir*+) ) | MG1655 RP4-2-Tc::[ΔMu1::aac(3)IV-ΔaphA-Δnic35-ΔMu2::zeo] ΔdapA::(erm-*pir*) ΔrecA |
| *E. coli* DH5α | F<sup>-</sup> *endA1 glnV44 thi-1 recA1 relA1 gyrA96 deoR nupG purB20* ϕ80dlacZΔM15 Δ(*lacZYA-argF*)U169, hsdR17(*rK<sup>-</sup>mK<sup>+</sup>*), λ<sup>-</sup>. For routine cloning. |
| Plasmid | Notes |
| pARO190 | Amp<sup>r</sup>; *bom*, basis of mobilization gens; oriT from RP4. *lacZα*, for blue-white screening (from ATCC) |
| pARO190-*rhsD*<sub>1616-1049</sub> | Contains part of *rhsD*, inserted from 3' to 5' |
| pARO190-*pspF-M* | 1<sup>st</sup> codon of modified from LEU (TTG) to MET (ATG). |
| pARO190-*PAS*<sub>(Δ2-304)</sub> |  |
| pARO190-*PAS*<sub>(Δ2-320)</sub> |  |
| pARO190-*GlrR*<sub>(Δ2-122)</sub> |  |
| pARO190-*GlrR*<sub>(Δ2-131)</sub> |  |
| pARO190-*tyrR*<sub>(Δ2-123)</sub> |  |
| pARO190-*tyrR*<sub>(Δ2-200)</sub> |  |
| pARO190-*prpR*<sub>(Δ2-208)</sub> | Could not be conjugated into *P. laumondii* TTO1. |

### Identification of putative σ^54^ binding sites

Putative sigma factor binding sites were located with the online Motif Locator (https://www.cmbl.uga.edu/software/motloc.html) **(Mrázek *et al*., 2008)**. *P. laumondii* TTO1 was selected from the list of previously uploaded genomes, while a list of validated intragenic and intergenic σ^54^ sites from *E. coli* MG1655 were used as input sequences **(Bonocora *et al*., 2015)**. Trial runs with the genome of *E. coli* MG1655 were used to determine the minimum motif score. A score of 14 best recapitulated the previous *E. coli* experiments (177 hits with Motif Locator vs 135 from ChIP-seq analysis; 75/135 identical in both).

Hypergeometric calculations were performed via the phyper package in R or with an online hypergeometric calculator (https://aetherhub.com/Apps/HyperGeometric). The coding regions of NRPS from the *P. laumondii* genome were added and used as the number of successes in the sample (228,311 of 5,612,110 bases). Results are reported as 1 – the cumulative distributive function, ie. the likelihood of discovering N or greater successes in the sample. Binomial tests were calculated in Excel, assuming a 50% likelihood of sense/antisense orientation.

### Metabolomics analysis

An overnight culture of *P. laumondii* containing each plasmid of interest was diluted 1:100 into tryptic soy broth supplemented with 100 µg/L carbenicillin, then incubated at 30 °C with shaking at 250 rpm for four days. 0.5 mM IPTG was added to induce bEBP production, while 2% v/v Amberlite XAD16 resin was used to capture produced metabolites. At the end of this incubation the beads and cells were collected by centrifugation at 4 krpm for five minutes, washed three times with Tris-buffered saline (pH 7.6), and then extracted with a 1:1 mixture of methanol:acetone. The solvent was then removed via speed-vac, and the residue was suspended in 1 mL MeOH. Leucine encephalin acetate was added to a final concentration of 1 μM to serve as an internal control. These extracts were then visualized on an Orbitrap LTQ Velos mass spectrometer (Thermo Electron Corporation, San Jose, CA) with a 2.7 μm Cortecs T3 reversed phase column (Waters Corporation, Milford, MA), measuring 2.1x 100 mm. 10 uL of each compound was injected, then separated at a constant flow rate of 0.3 mL/min, using a mixture of phase A (water containing 0.1% formic acid) and phase B (acetonitrile containing 0.1% formic acid) at the following gradient: 5% B for 1 min, a linear increase to 95% B at 20 min, isocratic with 95% B for 7 min, then a linear decrease to 5% B over 7 min. A full MS spectrum (m/z 150-2000) was acquired at a resolution of 100000, with the five most abundant singly and doubly charged ions were selected each second for fragmentation in a linear trap. Compound fragmentation was performed using a collision induced dissociation at normalized collision energy of 35%, with an activation time of 10 ms. The spectra were internally calibrated using diisooctyl phthalate (m/z 391.2843 Da) as a lock mass. All data was collected in triplicate, with three biological replicates for each bEBP.

### Data analysis

Compound Discoverer 3.3 and Thermo XCalibur Qual Browser were used to analyse the data collected from the LC-MS run. Compounds were selected based on a mass tolerance of 5ppm at MS1 and 0.6 Da at MS2, with compound identities tentatively assigned based on fragmentation patterns.

## Results and Discussion

### Selection of Photorhabdus as the test strain

*Photorhabdus* spp. are nematode-associated insect pathogens well-known for natural product biosynthesis (Chaston *et al*., 2011; Vizcaino *et al*., 2014; Bozhüyük *et al*., 2017). While attached to the gut of nematodes *Photorhabdus* spp. are commensal and express few if any natural products (Somvanshi *et al*., 2012). However, when the nematode infects an insect it will eject its less adhesive bacterial cargo. This variant of *Photorhabdus* spp. produces a wide variety of insecticidal proteins and antibiotics, killing the insect and sanitizing its corpse. This creates a nutrient-rich milieu that is ideal for the growth of both bacteria and nematodes (Chaston *et al*., 2011). Analysis of publicly available *Photorhabdus* genomes by antiSMASH (Weber *et al*., 2015) revealed two distinct clades (Fig. 1). The first contained four or fewer BGCs, in line with the majority of other sequenced Gammaproteobacteria (Fig. S1). The second clade was far more productive, with on average more than twenty BGCs (Fig. 1). Both clades were present in the three *Photorhabdus* subspecies (*P. asymbiotica; P. luminescens and P. laumondii; P. temperata*), suggesting that the divide is the result of evolutionary pressures and not an artifact of the small number of sequenced strains. As the genomes of *Photorhabdus* spp. rich in BGC also contain a large number of mobile genetic elements (Ogier *et al*., 2010), it seems likely that these bacteria have recently acquired a host of BGC to aid in insect pathogenesis. This includes *P. laumondii* TTO1, which contained 21 putative biosynthetic gene clusters as identified by antiSMASH (Table S2).

**Figure 1.**
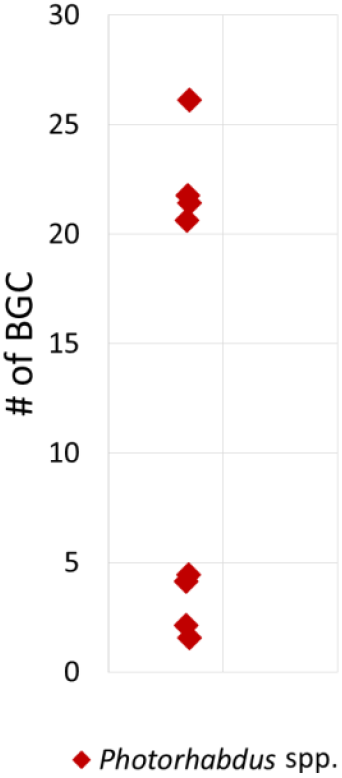
Number of biosynthetic gene clusters within *Photorhabdus* spp. This subset of publically available strains reveals two distinct *Photorhabdus* clades: those with a large number of biosynthetic gene clusters and those with significantly fewer. Genomes were obtained from NCBI and analyzed via antiSMASH 3.0. For a full list of the strains investigated see Table S6.

**Figure 2.**
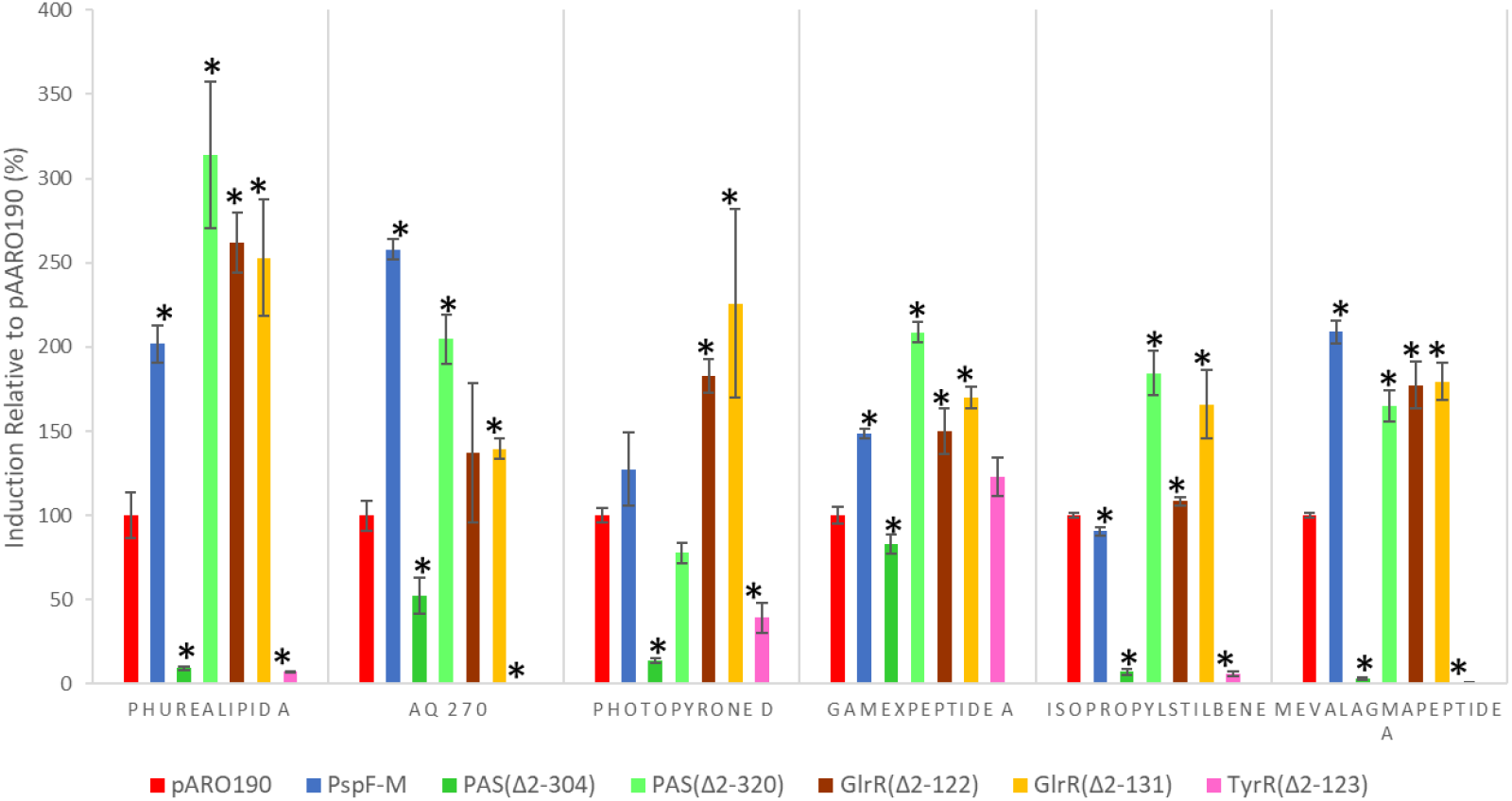
Metabolome following induction with bEBPs. Production of putative *P. laumondii* metabolites was scaled to pARO190. Intensities gathered by LCMS (positive mode), with compounds identities tentatively assigned based on MS/MS analysis. Stars indicate statistically significant variation relative to pARO190 (p < 0.05, two-tailed student T test).

A pBLAST search of the *P. laumondii* TTO1 genome with the *E. coli* variant of σ^54^ returned one hit with 77% identity and 88% similarity, putatively identified as a multispecies σ^54^. We then used the online bioinformatics tool motif locator to identify putative σ^54^ binding sites in the *P. laumondii* TTO1 genome (Mrázek *et al*., 2008; Bonocora *et al*., 2015) (Tables S3 and S4). A large number of such sites were found, in line with previous analyses of *E. coli* and *Salmonella enterica* genomes (Samuels *et al*., 2013; Bonocora *et al*., 2015). 89% of these sites were intragenic (211/236), well above the 62% identified in *E. coli* (Bonocora *et al*., 2015). Non-ribosomal peptide synthetase (NRPS) BGCs were particularly enriched in putative σ^54^ sites, with 20/236 such sites located within NRPS or NRPS-PKS coding regions, including in clusters known to produce the gameXPeptides and mevalagmapeptides (Tables 2, S3, and S4). This is significantly more than expected from a random distribution (Hypergeometric test *p* = 0.00175), suggesting σ^54^ plays a key role in the regulation of these biosynthetic pathways. As with previous reports we found that far more intragenic σ^54^ sites were in the sense orientation than would be expected due to chance (145/211, Binomial test *p* = 2.78 × 10^−8^). However, no similar enrichment was observed in sites within NRPS or NRPS-PKS (12/20, Binomial test *p* = 0.251).

**Table 2.** σ^54^ binding sites in *Photorhabdus laumondii* TTO1 which lie within NRPS or NRPS-PKS.

| BGC | Position | Gene | Orientation | Distance from gene start | Distance from gene end | Products | Reference |
| --- | --- | --- | --- | --- | --- | --- | --- |
| 5 | 1026921 | plu0898 | Antisense | 2924 | 1793 | Mevalagmapeptides | (Wang *et al.*, 2019) |
| 6 | 1280255 | plu1113 | Sense | 2985 | 11962 | Unknown |  |
| 6 | 1283423 | plu1113 | Sense | 6153 | 8794 | Unknown |  |
| 6 | 1286636 | plu1113 | Sense | 9366 | 5581 | Unknown |  |
| 6 | 1294572 | plu1115 | Sense | 945 | 4414 | Unknown |  |
| 7 | 1413798 | plu1220 | Sense | 3108 | 69 | Unknown |  |
| 8 | 2223806 | plu1878 | Antisense | 1416 | 1778 | Luminmycin A, gliobactin A, cepafungin | (Bian *et al.*, 2012; Wang *et al.*, 2019) |
| 8 | 2235424 | plu1880 | Sense | 8908 | 3558 | Luminmycin A, gliobactin A, cepafungin | (Bian *et al.*, 2012; Wang *et al.*, 2019) |
| 10 | 2724351 | plu2320 | Sense | 4076 | 2068 | Piscibactins | (Wang *et al.*, 2019) |
| 10 | 2724766 | plu2320 | Antisense | 1653 | 4491 | Piscibactins | (Wang *et al.*, 2019) |
| 10 | 2725347 | plu2320 | Sense | 5072 | 1072 | Piscibactins | (Wang *et al.*, 2019) |
| 10 | 2732439 | plu2321 | Antisense | 5706 | 6004 | Piscibactins | (Wang *et al.*, 2019) |
| 12 | 3143522 | plu2670 | Antisense | 30148 | 19017 | Kolossin | (Bode *et al.*, 2015) |
| 12 | 3144847 | plu2670 | Antisense | 28823 | 20342 | Kolossin | (Bode *et al.*, 2015) |
| 12 | 3148383 | plu2670 | Sense | 23878 | 25287 | Kolossin | (Bode *et al.*, 2015) |
| 12 | 3157460 | plu2670 | Antisense | 16210 | 32955 | Kolossin | (Bode *et al.*, 2015) |
| 14 | 3683221 | plu3130 | Sense | 4693 | 5241 | Unknown |  |
| **15** | 3873015 | plu3263 | Sense | 7923 | 7759 | GameXPeptides | (Nollmann<br>*et al.*, 2015) |
| **17** | 4147924 | plu3534 | Antisense | 7149 | 2647 | Unknown |  |
| **17** | 4156839 | plu3535 | Sense | 1750 | 8091 | Unknown |  |

### Plasmid construction and metabolomics

A pBLAST search of the *P. laumondii* TTO1 genome with the core domain of *dctD* revealed six homologues with an E values less than 1 × 10^−10^ (Table S5). Several of these had significant homology to better-characterized *E. coli* bEBPs, including *glnG, glrR, prpR, pspF*, and *tyrR*. Putative regulatory roles for these bEBPs include nitrogen regulation (*glnG*), propionate catabolism (*prpR*), phage shock (*pspF*), and tyrosine biosynthesis (*tyrR*), among others (Pittard and Davidson, 1991; Shiau *et al*., 1993; Jovanovic *et al*., 1996; Lee *et al*., 2005; Reichenbach *et al*., 2009). Of these five regulators, four are known to bind to σ54 in *E. coli*, while TyrR, despite its homology, instead interacts with σ^70^ (Camakaris *et al*., 2021). The last bEBP in *P. laumondii* had no significant homology to any *E. coli* bEBPs, but did contain a PAS regulatory domain. These domains have been previously found to bind to small molecules, indicating a potential chemosensory role (Liu *et al*., 2015). To determine which, if any, of the native *P. laumondii* bEBPs regulated natural product biosynthesis we then set out to create constitutively active variants of these proteins.

The domain structure of most bEBPs is well understood, with a central ATPase domain flanked on one side by a DNA recognition element, which binds to DNA 60-300 bp upstream of the σ^54^ promoter (Bush and Dixon, 2012). The other end of the bEBP generally contains a regulatory region, which ties bEBP activity to either a cognate kinase or the presence of a chemical signal (for example, NO for the NorR receptor) (Bush and Dixon, 2012). Truncation of the regulatory and DNA-binding regions can create constitutively active bEBPs like tDctD (Xu *et al*., 2004), while truncation of just the regulatory domain should in principle render the bEBP constitutively active but targeted only to its native genes. As the precise borders of the regulatory domains were not known, we prepared a number of truncated proteins, with cuts either at the end of the regulatory domain (as listed in the UniProt reference sequence), the beginning of the AAA+ ATPase domain, or midway between. In *E. coli* PspF is regulated by binding to an inhibitory protein, PspA, and lacks a regulatory domain. The *P. laumondii* homologue further had a mutated start codon, and so to address both issues we expressed it without truncation, with the start codon mutated to methionine. This allowed us to prepare five of the six *P. laumondii* bEBPs (Table 2), with four of these successfully conjugated into *P. laumondii*. Repeated attempts to conjugate pARO190-prpR_(Δ2-208)_ resulted in no colonies, indicating this bEBP may be toxic to *P. laumondii*.

Expression of the bEBPs drastically altered natural product biosynthesis in *P. laumondii*. Expression of a truncated copy of GlrR, GlrR_(Δ2-122)_, activated all six of the natural products under investigation, with an average increase in signal of 190% relative to pARO190 alone. The PAS-containing bEBP displayed differential activity based on its truncation point, with the variant truncated midway between the PAS and ATPase domains, PAS_(Δ2-304)_, broadly inactivating the compounds under investigation and the variant truncated to the beginning of the ATPase domain activating five of the six compounds. While more work will be required to determine why PAS_(Δ2-304)_ suppresses natural product production, this behaviour would be expected if an inactive ATPase domain were to bind to and prevent the activation of σ^54^. TyrR_(Δ2-123)_ also generally suppressed natural product production, though like PAS_(Δ2-304)_ it had little effect on the production of gameXPeptide A. Further truncation of TyrR led to TyrR_(Δ2-200)_. This protein suppressed production of all six natural below detectable limits, with cells forming exclusively small, yellow colonies when grown on agar. These features are consistent with the mutualistic form of *P. laumondii*, suggesting that TyrR may play a role in growth of the bacteria when in its nematodal host (Somvanshi *et al*., 2012).

While this induction was impressive, comparison between pARO190 and plasmid-free *P. laumondii* revealed that even in the absence of bEBP significant induction was occurring. All six compounds increased in titre when pARO190 was present, with IPS in particular increasing 985% (Fig. 3). Built from fusing the pUC19 backbone with a fragment of pSUP2021 (Parke, 1990), pARO190 contains both *lacZ* in the multiple cloning site (for blue-white screening) and several components of the *tra* conjugation machinery. At the moment it is not clear which, if any, of these underlies the induction effect we observe here.

**Figure 3.**
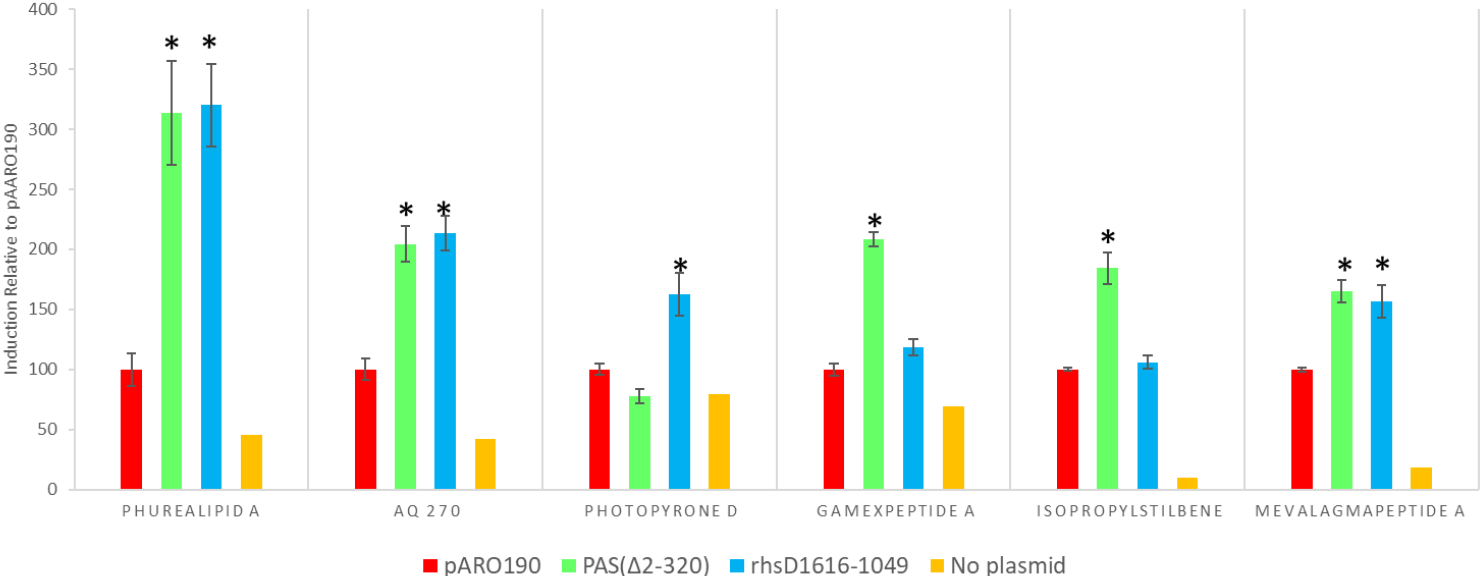
Production in the absence of plasmid and with pARO190-*rhsD*_*1616-1049*_. Production was scaled to pARO190, with PAS included for context. Intensities gathered by LCMS (positive mode), with compounds identities tentatively assigned based on MS/MS analysis. Stars indicate statistically significant variation relative to pARO190 (p < 0.05, two-tailed student T test).

During data analysis we also discovered that one of the plasmids we had analyzed did not contain a bEBP. Originally intended to be a constitutively active variant of DctD from *S. melilloti*, due to a labeling error pARO190 was instead loaded with a fragment of *E. coli rhsD*. The gene was inserted into the multiple cloning site in a 3′ to 5′ direction. Largely non-coding, the first open reading from of pARO190-*rhsD*_1616-1049_ is peptide only 15 residues long. While unlikely to encode a transcription factor, expression of pARO190-*rhsD*_1616-1049_ was roughly as activating as the best of the bEBPs tested (Fig 3). Production of compounds with MS fragmentation patterns consistent with phurealipid A and AQ-270 more than doubled relative to pARO190 (320% and 214%, respectively). Statistically significant increases were also seen in peaks consistent with photopyrone D (163%), gameXPeptide A (118%), and mevalagmapeptide A (156%). Of the compounds tested, only isopropylstilbene was unaffected. Given the broad effect of this sequence, and the general activation observed with pARO190, it seems likely that the induction we see here is not due to specific interactions by the bEBPs, but rather some form of generalized stress response, triggered by either plasmid carriage or protein synthesis.

## Conclusion

Natural product discovery is limited by an inability to effectively control *in situ* natural product biosynthesis. By conjugating a number of plasmids into *P. laumondii* TTO1 we have been able to broadly induce biosynthesis in this strain. The varied levels of induction and lack of a unifying feature in the cargo of our induction plasmids suggest that the increase in biosynthesis is not due to any specific transcription factor, but rather due to the stress imposed by pARO190 and its cargo. This suggests a genome-agnostic way of activating silent biosynthetic gene clusters in this species.

## Author Contributions

LR cloned the pARO190-bEBP plasmids and carried out the LCMS analysis. MS sequenced pARO190-*rhsD*_1616-1049_. MA conducted early experiments with tdctD. BF conceived of and designed the experiments, supervised the work, and performed the bioinformatics analysis.

## Acknowledgements

We are indebted to Lisa Freeman, Heng Jiang, and Marcos Di Falco for helpful comments and technical support, and to the Concordia Centres for Biological Applications of Mass Spectrometry and Structural and Functional Genomics for access to necessary equipment. This work was supported by the Natural Sciences and Engineering Research Council of Canada (NSERC).

## Additional Information

**Supplementary information** accompanies this paper.

