## Supplementary Information for "Genes acquired by horizontal gene transfer share common regulatory patterns in *Photorhabdus laumondii*"

Lydia Rili[1], Marcus Simoes[1], Maythem Ali[1], Brandon L. Findlay[1,2,*]

Affiliations:

[1] Department of Chemistry and Biochemistry, Concordia University, Montreal, Québec, Canada

[2] Department of Biology, Concordia University, Montreal, Québec, Canada

[*] Corresponding author.

Brandon L. Findlay,

Sequence of the synthesized dctD$_{141\text{-}394}$ gBlock. Flanking KpnI/HindIII restriction digest sites have been added to aid in cloning.

GGTACCATGGTCGTCGCCACGCTGCTGCACCAATGGAGCCGCCGCAGGACCGGCAATTTCGTCGCGCTGAATTGCGGC
GCTCTGCCGGAAACGGTGATCGAAAGCGAGCTCTTCGGCCACGAGCCCGGCGCCTTTACCGGCGCCGTCAAGAAGCGG
ATCGGCCGGATCGAGCATGCGAGCGGCGGCACGCTCTTCCTCGACGAGATCGAGGCCATGCCGCCGGCAACTCAGGTG

AAGATGCTGCGCGTGCTGGAAGCCCGCGAGATCACGCCGCTCGGCACCAATCTGACCCGCCCCGTCGACATCCGCGTC
GTCGCCGCAGCCAAGGTCGATCTCGGCGACCCGGCCGCGCGCGGCGATTTCCGCGAGGATCTCTATTACCGGCTGAAC
GTCGTGACGCTCTCGATCCCGCCCCTGCGCGAACGGCGCGACGACATCCCCCTCCTCTTCTCCCATTTCCTGGCCCGCGC
CTCGGAACGCTTCGGCCGCGAAGTGCCCGCGATCTCGGCTGCCATGCGCGCGTACCTGGCGACGCATTCCTGGCCCGGC
AATGTGCGCGAGCTTTCGCACTTCGCCGAACGGGTGGCGCTGGGGGTGGAGGGAAACCTGGGAGTTCCGGCCGCAGCG
CCCGCCTCAAGCGGAGCGACCCTGAAGCTT

**Figure S1. antiSMASH analysis of Gammaproteobacteria.** Strains were selected from publically available databases, then analyzed under default antiSMASH settings. *Streptomyces* spp. included for scale.  The strains are listed in table S4.

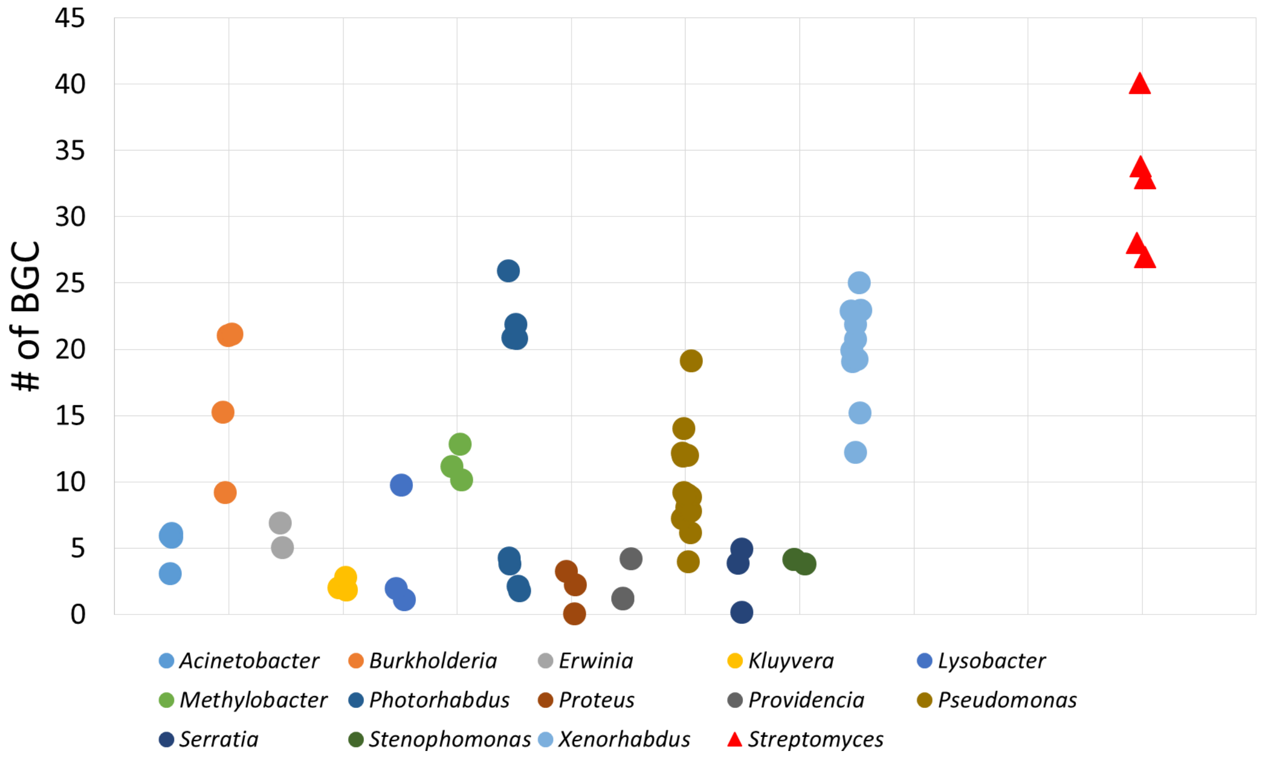

**Table S1.** Primers for amplification of EBPs from *Photorhabdus laumondii* TTO1.

| **Primers** | | |
|---|---|---|
| FW-*dctD*$_{(141\text{-}394)}$ | RE HindIII; ATG start codon | CGCGCAAGCTTCCGG**ATG**GAAGGCCTGCCGCT |
| RC-*dctD*$_{(141\text{-}394)}$ | RE KpnI; TAA stop codon; 6xHis tag | CCGCCGGTACCGCGCG**TTA**GTGGTGGTGGTGGTGGTGCAGGGTCGCTCCGC |
| FW-*pspF-M* | RE HindIII; ATG start codon | CGCGCAAGCTTCCGG**ATG**TCGGTGAAAATAACCATG |
| RC-*pspF* | RE KpnI; TAA stop codon; 6xHis tag | CCGCCGGTACCGCGCG**TTA**GTGGTGGTGGTGGTGGTGCTTCTCGCCGACGTTG |
| FW-*uPAS*$_{(\Delta 2\text{-}304)}$ | RE HindIII; ATG start codon | CGCGCAAGCTTCCGG**ATG**GAACAGCAAAATGAATATCT |
| FW-*uPAS*$_{(\Delta 2\text{-}320)}$ | RE HindIII; ATG start codon | CGCGCAAGCTTCCGG**ATG**TATGATGAAATTATTGGCAGA |
| RC-*uPAS* | RE KpnI; TAA stop codon; 6xHis tag | CCGCCGGTACCGCG**TTA**GTGGTGGTGGTGGTGGTGATAATCAACTCCTAATTTCTTTAAG |
| FW-*GlrR*$_{(\Delta 2\text{-}122)}$ | RE HindIII; ATG start codon | CGCGCAAGCTTCCGG**ATG**GCGTTGACCACGCC |
| FW-*GlrR*$_{(\Delta 2\text{-}131)}$ | RE HindIII; ATG start codon | CGCGCAAGCTTCCGG**ATG**CAGTGGCGGGAACAG |
| RC-*GlrR* | RE SalI; TAA stop codon; 6xHis tag | CCGCCGTCGACGCGCG**TTA**GTGGTGGTGGTGGTGGTGTTCTTTAAAATCATTCGCATCC |
| FW-*tyrR*$_{(\Delta 2\text{-}123)}$ | RE HindIII; ATG start codon | CGCGCAAGCTTCCGG**ATG**CCTATCGGGCAGTTTATCAG |
| FW-*tyrR*$_{(\Delta 2\text{-}200)}$ | RE HindIII; ATG start codon | CGCGCAAGCTTCCGG**ATG**GGTAGTGAGTTTAAGGCGA |
| RC-*tyrR* | RE KpnI; TAA stop codon; 6xHis tag | CCGCCGGTACCGCGCG**TTA**GTGGTGGTGGTGGTGGTGTTCGTCACCATCCGGC |
| FW-*prpR*$_{(\Delta 2\text{-}208)}$ | RE HindIII; ATG start codon | CGCGCAAGCTTCCGG**ATG**TATCCAACCGAGAACAA |

| | | |
|---|---|---|
| RC-*prpR* | RE KpnI; TAA stop codon; 6xHis tag | CCGCCGGTACCGCGCG**TTA**GTG GTGGTGGTGGTGGTGTTCTCTT TCTCTTGCTATCTC |

**Table S2.** antiSMASH analysis of *Photorhabdus laumondii* TTO1.

| Cluster | Compound Class | Start | End | Most Similar BGC[a] |
|---|---|---|---|---|
| **1** | Bacteriocin | 72425 | 83348 | - |
| **2** | Blactam | 182436 | 203941 | - |
| **3** | Other | 294972 | 335676 | - |
| **4** | Bacteriocin | 579213 | 589992 | - |
| **5** | Nrps | 1001625 | 1055887 | Rhabdopeptides_biosynthetic_gene_cluster (100% of genes show similarity) |
| **6** | Nrps | 1257282 | 1319002 | Massetolide_A_biosynthetic_gene_cluster (100% of genes show similarity) |
| **7** | Nrps | 1373004 | 1433883 | - |
| **8** | Nrps-T1pks | 2202028 | 2258998 | Luminmycin_biosynthetic_gene_cluster (100% of genes show similarity) |
| **9** | Resorcinol | 2529713 | 2593940 | Indigoidine_biosynthetic_gene_cluster (60% of genes show similarity) |
| **10** | Nrps-T1pks | 2700286 | 2762991 | Yersiniabactin_biosynthetic_gene_cluster (5% of genes show similarity) |
| **11** | Other | 3072574 | 3115324 | - |
| **12** | Nrps | 3104571 | 3193674 | Streptomycin_biosynthetic_gene_cluster (2% of genes show similarity) |
| **13** | Nrps | 3215684 | 3265399 | Turnerbactin_biosynthetic_gene_cluster (23% of genes show similarity) |
| **14** | Nrps | 3626119 | 3716166 | Tolaasin_biosynthetic_gene_cluster (50% of genes show similarity) |
| **15** | Nrps | 3845127 | 3900777 | - |
| **16** | Phosphoglycolipid | 3983869 | 4005998 | - |
| **17** | Nrps | 4112061 | 4190703 | - |

| | | | | |
|---|---|---|---|---|
| **18** | Nrps | 4578801 | 4621714 | Zorbamycin_biosynthetic_gene_cluster (4% of genes show similarity) |
| **19** | T2pks | 4884466 | 4925752 | Anthraquinone_biosynthetic_gene_cluster (100% of genes show similarity) |
| **20** | Terpene | 5058951 | 5082523 | Carotenoid_biosynthetic_gene_cluster (83% of genes show similarity) |
| **21** | Siderophore | 5380534 | 5405194 | Desferrioxamine_B_biosynthetic_gene_cluster (60% of genes show similarity) |

[a]As determined by antiSMASH.

**Table S3.** $\sigma^{54}$ binding sites in *Photorhabdus laumondii* TTO1 which lie outside of gene coding regions.

| ID | Sequence | Position | Motif Strand | Upstream Gene | Gene Strand | Distance to Genes | Downstream Gene | Gene Strand | Notes[a] |
|---|---|---|---|---|---|---|---|---|---|
| **OS01** | TGCAACAGGCGTACCAT | 155112 | - | plu0148 | + | 113 ~~~~~ 123 | plu0149 | - | |
| **OS02** | TGCGAAAAGCATGCCAA | 251726 | - | plu0237 | - | 154 ~~~~~ 262 | plu0238 | + | |
| **OS03** | CTGGCATAGTTATTGCT | 393625 | + | plu0365 | - | -6 ~~~~~ -7 | plu0366 | - | |
| **OS04** | TTGGCACAATGCATGCT | 416684 | + | plu0385 | + | 141 ~~~~~ 318 | plu0386 | + | |
| **OS05** | TGCAATAAATATGCCCA | 917825 | - | plu0797 | - | 182 ~~~~~ 701 | plu0798 | - | |
| **OS06** | TTGGCAGTAACTTTGCA | 1300011 | + | plu1116 | - | 97 ~~~~~ -16 | plu1117 | + | Within BGC |
| **OS07** | AGCAAAAATCAGGCCAA | 1338456 | - | plu1150 | - | 141 ~~~~~ 639 | plu1151 | + | |
| **OS08** | TGCGAGAACTGTGCCAA | 1424272 | - | plu1232 | - | 63 ~~~~~ 171 | plu1233 | + | Within BGC |
| **OS09** | AGCAATTAGCATGCCAA | 1517040 | - | plu1307 | - | 285 ~~~~~ 251 | plu1308 | - | |
| **OS10** | AGCAAAAAACAGGCCAA | 1640545 | - | plu1368 | - | 141 ~~~~~ 1427 | plu1369 | + | |
| **OS11** | TGCAATATTAGTGCCTA | 1821961 | - | plu1519 | - | 180 ~~~~~ 503 | plu1520 | - | |
| **OS12** | TTGGCGTCTTTTTTGCT | 2081279 | + | plu1744 | + | 163 ~~~~~ 141 | plu1745 | + | |
| **OS13** | AGCAAAAATAGCACCAG | 2114529 | - | plu1771 | + | 4 ~~~~~ 105 | plu1772 | - | |
| **OS14** | ATCAAGAGTCGTGCCAA | 3027703 | - | plu2585 | - | 42 ~~~~~ 108 | plu2586 | + | |
| **OS15** | TTGGCAAGACCATTGCA | 3202741 | + | plu2700 | + | 287 ~~~~~ 53 | plu2701 | + | |
| **OS16** | ATGGCGCGGTTTTTGTA | 3920588 | + | plu3305 | - | 317 ~~~~~ 66 | plu3306 | - | |
| **OS17** | TTGGCGTAGACTATGCT | 3959532 | + | plu3329 | + | 199 ~~~~~ 197 | plu3330 | + | |
| **OS18** | AGCAACATACGTACCAA | 4174442 | - | plu3542 | - | 31 ~~~~~ 264 | plu3543 | + | Within BGC |

| | | | | | | | | |
|---|---|---|---|---|---|---|---|---|
| **OS19** | AGCAAAAACAATACCAC | 4330353 | - | plu3676 | + | 14 ~~~~~ 358 | plu3677 | + |
| **OS20** | AGCAAAAAACAGGCCAA | 4384994 | - | plu3719 | - | 141 ~~~~~ 652 | plu3720 | - |
| **OS21** | AGCAAAGGGTGTGCCAG | 4531475 | + | plu3857 | - | 30 ~~~~~ 121 | plu3858 | - |
| **OS22** | ATGGTACGCATATTGCT | 5024920 | + | plu4300 | - | 209 ~~~~~ 208 | plu4301 | + |
| **OS23** | TGCATTTAGTGTGCCAA | 5419988 | + | plu4649 | - | 593 ~~~~~ 379 | plut073 | - |
| **OS24** | TGCAAAATACATGCCAT | 5436009 | + | plu4662 | - | 321 ~~~~~ 34 | plu4663 | - |
| **OS25** | TTGGCGTTGTTTATGCT | 5487880 | + | plu4705 | - | 145 ~~~~~ 127 | plu4706 | - |

[a]Biosynthetic gene cluster boundaries as determined by antiSMASH.

**Table S4.** $\sigma^{54}$ binding sites in *Photorhabdus laumondii* TTO1 which lie inside of gene coding regions.

| **ID** | **Sequence** | **Position** | **Motif Strand** | **Gene** | **Gene Strand** | **Distance to Gene Ends** | **Notes[a]** |
|---|---|---|---|---|---|---|---|
| **IS01** | AGCAATTATTGTTCCAG | 32291 | - | plu0037 | - | 1368 ~~~~~ 512 | |
| **IS02** | CTGGCACCGTTCCTGCC | 46498 | + | plu0052 | + | 468 ~~~~~ 508 | |
| **IS03** | CTGGCATCATCATTGCG | 57114 | + | plu0061 | - | 1040 ~~~~~ 348 | |
| **IS04** | AGCAATTTTTCTGCCAT | 74760 | - | plu0076 | + | 397 ~~~~~ 271 | Within BGC |
| **IS05** | CCGGCACTGCTTTTGCC | 93324 | + | plu0097 | + | 660 ~~~~~ 1456 | |
| **IS06** | TTGGTGAGATTTTTGCA | 109714 | + | plu0108 | + | 910 ~~~~~ 2475 | |
| **IS07** | AGCAAAATTGTTGCCAC | 112672 | - | plu0109 | + | 470 ~~~~~ 710 | |
| **IS08** | GCGGCAAAGTTATTGCA | 154049 | + | plu0146 | + | 851 ~~~~~ 354 | |
| **IS09** | TGCAATATCACCGCCAA | 187947 | - | plu0176 | + | 493 ~~~~~ 915 | Within BGC |
| **IS10** | TGCATACAACGTGCCAT | 256195 | - | plu0240 | - | 670 ~~~~~ 781 | |
| **IS11** | AGCATCAGCCGTGCCAC | 309593 | - | plu0292 | - | 310 ~~~~~ 246 | |
| **IS12** | GTGGTACTGTTTTTGCT | 335238 | + | plu0316 | - | 1091 ~~~~~ 3372 | |
| **IS13** | GGCAATATCCGCGCCGA | 335828 | - | plu0316 | - | 1681 ~~~~~ 2782 | |
| **IS14** | CGCGATAACCGCGCCAG | 403257 | - | plu0374 | + | 396 ~~~~~ 430 | |
| **IS15** | AGCAAGGATCGCGCCGC | 409326 | - | plu0379 | - | 558 ~~~~~ 25 | |
| **IS16** | CTGGTACCGGTTATGCT | 472674 | + | plu0440 | + | 4087 ~~~~~ 129 | |
| **IS17** | TTGGCAGAATTCTTGCC | 480535 | + | plu0452 | + | 569 ~~~~~ 504 | |
| **IS18** | ATGGCGCGAACCATGCT | 489383 | + | plu0459 | + | 1122 ~~~~~ 430 | |
| **IS19** | GTGGTGTGATTTTTGCT | 490320 | + | plu0460 | + | 491 ~~~~~ 396 | |

| | | | | | | | |
|---|---|---|---|---|---|---|---|
| **IS20** | CCGGTGCGGCTTTTGCA | 506841 | + | plu0470 | - | 659 ~~~~~ 1741 | |
| **IS21** | TCGGCATAATTTTTGCC | 553575 | + | plu0504 | - | 790 ~~~~~ 297 | |
| **IS22** | ATGGTGTAGTTTTTGCT | 619418 | + | plu0549 | - | 841 ~~~~~ 834 | |
| **IS23** | CTGGTGTGGATGTTGCT | 638063 | + | plu0563 | + | 2038 ~~~~~ 420 | |
| **IS24** | GGCAATTATTCTGCCAG | 638707 | - | plu0564 | + | 205 ~~~~~ 708 | |
| **IS25** | AGCAAAATCAGTGCCAC | 648640 | - | plu0572 | + | 788 ~~~~~ 137 | |
| **IS26** | GTGGCGTTTTTTTTGCA | 750366 | + | plu0653 | + | 226 ~~~~~ 165 | |
| **IS27** | GCGGCGTGGTTTTTGCG | 754317 | + | plu0656 | + | 5 ~~~~~ 393 | |
| **IS28** | ATGGCACGACTGATGCC | 755111 | + | plu0657 | + | 338 ~~~~~ 1417 | |
| **IS29** | GTGGCACAACACTTGCC | 837773 | + | plu0729 | + | 58 ~~~~~ 180 | |
| **IS30** | GGCAAATTAAGTGCCAC | 841100 | - | plu0733 | + | 181 ~~~~~ 21 | |
| **IS31** | TGCATTATCCGTACCAC | 841724 | - | plu0734 | - | 519 ~~~~~ 592 | |
| **IS32** | TGCATTATCGGTGCCAC | 841868 | - | plu0734 | - | 663 ~~~~~ 448 | |
| **IS33** | CGCATTATCGGTGCCAC | 842009 | - | plu0734 | - | 804 ~~~~~ 307 | |
| **IS34** | CGCATTATCTGTGCCAC | 843578 | - | plu0735 | - | 798 ~~~~~ 316 | |
| **IS35** | AGCATATGCAGTGCCAG | 851499 | - | plu0740 | - | 70 ~~~~~ 927 | |
| **IS36** | TGCATTTTCTGTGCCGG | 914043 | - | plu0794 | - | 493 ~~~~~ 762 | |
| **IS37** | GGCATTATTTACGCCAG | 937571 | - | plu0806 | + | 3984 ~~~~~ 429 | |
| **IS38** | TTGGTACGGCATTTGCC | 968434 | + | plu0836 | + | 243 ~~~~~ 552 | |
| **IS39** | TGCATTTTGCATACCAA | 1026921 | - | plu0898 | + | 1793 ~~~~~ 2924 | Within BGC |
| **IS40** | ATGGTACAATTATTGCG | 1076598 | + | plu0935 | + | 175 ~~~~~ 942 | |
| **IS41** | CTGGTGCGGATATTGCG | 1094444 | + | plu0954 | + | 174 ~~~~~ 24 | |
| **IS42** | GCGGCGGGGATATTGCA | 1104574 | + | plu0961 | - | 1545 ~~~~~ 2879 | |
| **IS43** | CTGGCGTGTAAATTGCG | 1114695 | + | plu0962 | - | 7183 ~~~~~ 391 | |
| **IS44** | TGCATATATTGTGCCCC | 1139301 | - | plu0970 | - | 2743 ~~~~~ 4588 | |
| **IS45** | CCGGCACGAAACCTGCT | 1165201 | + | plu0981 | - | 950 ~~~~~ 422 | |
| **IS46** | CTGGCACCGCTTTCGCT | 1178843 | + | plu0995 | - | 475 ~~~~~ 2022 | |
| **IS47** | TTGGCGCGTTATTTGCC | 1280255 | + | plu1113 | + | 2985 ~~~~~ 11962 | Within BGC |
| **IS48** | TTGGCCCGCTACTTGCC | 1283423 | + | plu1113 | + | 6153 ~~~~~ 8794 | Within BGC |
| **IS49** | TTGGCGCGTTATTTGCC | 1286636 | + | plu1113 | + | 9366 ~~~~~ 5581 | Within BGC |

| | | | | | | | |
|---|---|---|---|---|---|---|---|
| **IS50** | TTGGCGCGCTACTTGCC | 1294572 | + | plu1115 | + | 945 ~~~~~ 4414 | Within BGC |
| **IS51** | TGCAGGACTCGTGCCTG | 1376361 | - | plu1192 | - | 1873 ~~~~~ 559 | Within BGC |
| **IS52** | TTGGCGTGATTGCTGCA | 1384072 | + | plu1200 | + | 250 ~~~~~ 330 | Within BGC |
| **IS53** | ATGGCACGGGATTCGCT | 1413798 | + | plu1220 | + | 3108 ~~~~~ 69 | Within BGC |
| **IS54** | TGCATAAGTTGTGCCGT | 1438818 | - | plu1244 | + | 623 ~~~~~ 624 | |
| **IS55** | CTGGCCTGCATATTGCC | 1460313 | + | plu1262 | + | 364 ~~~~~ 708 | |
| **IS56** | GGCAAAATCCATGCCTG | 1484495 | - | plu1275 | - | 459 ~~~~~ 1077 | |
| **IS57** | CCGGCGTAGATGATGCA | 1488576 | + | plu1279 | - | 574 ~~~~~ 168 | |
| **IS58** | GGCAAGTTCCATACCAA | 1493396 | - | plu1283 | - | 798 ~~~~~ 402 | |
| **IS59** | ATGGCGTGATATATGCA | 1659119 | + | plu1379 | + | 772 ~~~~~ 399 | |
| **IS60** | CTGGAATGAAACCTGCA | 1674903 | + | plu1395 | - | 302 ~~~~~ 708 | |
| **IS61** | CTGGTGCGATTATTGCG | 1723973 | + | plu1433 | + | 742 ~~~~~ 117 | |
| **IS62** | AGCATTACCTATGCCAA | 1730064 | - | plu1438 | + | 227 ~~~~~ 530 | |
| **IS63** | GTGGCGCTGTTATTGCG | 1735983 | + | plu1442 | + | 793 ~~~~~ 396 | |
| **IS64** | CTGGTGCGGGTAATGCA | 1747312 | + | plu1455 | + | 145 ~~~~~ 1143 | |
| **IS65** | CTGGCTTCAAATTTGCA | 1765108 | + | plu1471 | - | 389 ~~~~~ 432 | |
| **IS66** | TTGGCATCAGCTTTGCA | 1771229 | + | plu1477 | + | 253 ~~~~~ 429 | |
| **IS67** | AGCAATATCTTCGCCAG | 1843091 | - | plu1540 | - | 804 ~~~~~ 433 | |
| **IS68** | TCGGCATTGGTTTTGCT | 1845786 | + | plu1542 | + | 790 ~~~~~ 366 | |
| **IS69** | GTGGCACCAATTCTGCA | 1865009 | + | plu1560 | + | 464 ~~~~~ 937 | |
| **IS70** | TGCAATAATGGTACCGC | 1895401 | - | plu1589 | - | 384 ~~~~~ 511 | |
| **IS71** | TTGGCGCAATTTATGCT | 1915850 | + | plu1604 | + | 534 ~~~~~ 782 | |
| **IS72** | CTGGCACTGTTAATGCT | 1986385 | + | plu1661 | - | 520 ~~~~~ 147 | |
| **IS73** | GGCATAAAGCATGCCAA | 2022435 | - | plu1694 | - | 640 ~~~~~ 906 | |
| **IS74** | CTGGCGCTTCATTTGCA | 2043765 | + | plu1713 | - | 452 ~~~~~ 387 | |
| **IS75** | GTGGCACAATATATGCC | 2062778 | + | plu1728 | - | 514 ~~~~~ 871 | |
| **IS76** | GGCAAAATTCAGGCCAT | 2088755 | - | plu1751 | - | 724 ~~~~~ 511 | |
| **IS77** | TTGGCGTGCTGATTGCT | 2196984 | + | plu1849 | + | 602 ~~~~~ 279 | |
| **IS78** | CCGGCATGGCAATTGCT | 2223806 | + | plu1878 | - | 1778 ~~~~~ 1416 | Within BGC |
| **IS79** | CGCAATTTCGATGCCAG | 2235424 | - | plu1880 | - | 8908 ~~~~~ 3558 | Within BGC |

| | | | | | | | |
|---|---|---|---|---|---|---|---|
| **IS80** | AGCAACACCCCCGCCAG | 2306273 | - | plu1939 | - | 310 ~~~~~ 34 | |
| **IS81** | GGCACAACTCATGCCAG | 2311693 | - | plu1946 | - | 26 ~~~~~ 662 | |
| **IS82** | GGCAATTTCTGTACCAG | 2329959 | - | plu1959 | - | 1290 ~~~~~ 205 | |
| **IS83** | CTGGCACGTTATCTGCT | 2365730 | + | plu1994 | + | 45 ~~~~~ 844 | |
| **IS84** | GGCAAATAGAATGCCAG | 2396887 | - | plu2026 | - | 469 ~~~~~ 63 | |
| **IS85** | AGCAATTATCAGGCCAA | 2412641 | - | plu2042 | - | 498 ~~~~~ 493 | |
| **IS86** | TTGGCACAGTTGTTGTA | 2505524 | + | plu2120 | - | 614 ~~~~~ 285 | |
| **IS87** | CTGGTGCGCTTCCTGCA | 2512172 | + | plu2128 | - | 74 ~~~~~ 268 | |
| **IS88** | AGCATAAAGTATGCCAA | 2524888 | - | plu2142 | + | 4 ~~~~~ 635 | |
| **IS89** | GGCAAGTTTGACGCCAG | 2537441 | - | plu2152 | + | 373 ~~~~~ 367 | Within BGC |
| **IS90** | GGCAAGTTTGACGCCAG | 2537507 | - | plu2152 | + | 439 ~~~~~ 301 | Within BGC |
| **IS91** | GGCAAGTTTGACGCCAG | 2537573 | - | plu2152 | + | 505 ~~~~~ 235 | Within BGC |
| **IS92** | ATGGTATGGCTGTTGCA | 2555218 | + | plu2168 | + | 955 ~~~~~ 429 | Within BGC |
| **IS93** | TTGGCGCGATTGTTGCT | 2567780 | + | plu2183 | + | 130 ~~~~~ 330 | Within BGC |
| **IS94** | TTGGCGTGATTATTGCG | 2619582 | + | plu2226 | + | 679 ~~~~~ 528 | |
| **IS95** | TTGGCACAATTTTTGTT | 2673205 | + | plu2275 | + | 154 ~~~~~ 1160 | |
| **IS96** | CTGGCGCAAAAAATGCC | 2684509 | + | plu2286 | - | 1087 ~~~~~ 156 | |
| **IS97** | GGCAAAATTTCTGCCAG | 2685908 | - | plu2287 | - | 1177 ~~~~~ 1389 | |
| **IS98** | TGCGTGTATTGTGCCAG | 2686784 | - | plu2287 | - | 2053 ~~~~~ 513 | |
| **IS99** | GTGGCACAAATTGTGCA | 2696327 | + | plu2295 | - | 758 ~~~~~ 1248 | |
| **IS100** | CCGGTGTGGATTTTGCA | 2716511 | + | plu2317 | + | 1051 ~~~~~ 177 | |
| **IS101** | CTGGCACAACATCTGCC | 2724351 | + | plu2320 | + | 4076 ~~~~~ 2068 | Within BGC |
| **IS102** | AGCAAAATGAGTACCAG | 2724766 | - | plu2320 | + | 4491 ~~~~~ 1653 | Within BGC |
| **IS103** | CTGGCACCACAATTGCC | 2725347 | + | plu2320 | + | 5072 ~~~~~ 1072 | Within BGC |
| **IS104** | TGCAAATTTGCTGCCAG | 2732439 | - | plu2321 | + | 6004 ~~~~~ 5706 | Within BGC |
| **IS105** | TTGGCATAGAAAATGCG | 2754201 | + | plu2337 | + | 270 ~~~~~ 140 | Within BGC |
| **IS106** | ATGGCACTGAAAATGCA | 2809605 | + | plu2395 | + | 1108 ~~~~~ 2940 | |
| **IS107** | TTGGCGTCGATATTGCA | 2873222 | + | plu2445 | + | 30 ~~~~~ 516 | |
| **IS108** | AGCAAAATCAATACCAT | 2923724 | - | plu2482 | - | 1168 ~~~~~ 589 | |
| **IS109** | TTGGCGCGTTTATTGCG | 2935234 | + | plu2494 | - | 520 ~~~~~ 1113 | |

| | | | | | | | |
|---|---|---|---|---|---|---|---|
| **IS110** | GGCAACTTCTGCGCCAT | 2936196 | - | plu2494 | - | 1482 ~~~~~ 151 | |
| **IS111** | CGGGCGCTGTTTTTGCT | 2945393 | + | plu2501 | - | 212 ~~~~~ 776 | |
| **IS112** | CGCAAATACCCTGCCAG | 2947509 | - | plu2502 | - | 943 ~~~~~ 78 | |
| **IS113** | CCGGTATACATATTGCT | 2951607 | + | plu2507 | + | 567 ~~~~~ 336 | |
| **IS114** | TGCATTAATTGTGCCAG | 3025365 | - | plu2582 | - | 466 ~~~~~ 924 | |
| **IS115** | AGCAATTTTTGTGCCTC | 3064881 | - | plu2621 | - | 31 ~~~~~ 1449 | |
| **IS116** | AGCAAGACCTACGCCAT | 3077228 | - | plu2628 | - | 846 ~~~~~ 1528 | Within BGC |
| **IS117** | GTGGTGCAACTTTTGCA | 3082085 | + | plu2632 | + | 597 ~~~~~ 1422 | Within BGC |
| **IS118** | CTGGCGCTACACTTGCA | 3090895 | + | plu2640 | - | 248 ~~~~~ 206 | Within BGC |
| **IS119** | CTGGCCTGCTATCTGCA | 3092835 | + | plu2642 | + | 261 ~~~~~ 2473 | Within BGC |
| **IS120** | TTGGCGCGGCAGTTGCA | 3143522 | + | plu2670 | - | 19017 ~~~~~ 30148 | Within BGC |
| **IS121** | CCGGCACCGGTAATGCT | 3144847 | + | plu2670 | - | 20342 ~~~~~ 28823 | Within BGC |
| **IS122** | GGCAAATAGCGGGCCAA | 3148383 | - | plu2670 | - | 23878 ~~~~~ 25287 | Within BGC |
| **IS123** | TTGGAATAGATCATGCA | 3157460 | + | plu2670 | - | 32955 ~~~~~ 16210 | Within BGC |
| **IS124** | CGCAAAAGTCGCGCCAA | 3232845 | - | plu2725 | - | 1495 ~~~~~ 537 | Within BGC |
| **IS125** | TTGGCATAAATTCTGCT | 3285107 | + | plu2771 | - | 602 ~~~~~ 44 | |
| **IS126** | CGCAAAAACTGAGCCAG | 3384954 | - | plu2850 | - | 1990 ~~~~~ 55 | |
| **IS127** | TTGGCCCAACAAATGCA | 3386183 | + | plu2852 | - | 230 ~~~~~ 740 | |
| **IS128** | CGCAAAAATTGGGCCAA | 3388848 | - | plu2854 | - | 264 ~~~~~ 1279 | |
| **IS129** | AGCAACTCCAGTGCCAG | 3462937 | - | plu2959 | - | 99 ~~~~~ 483 | |
| **IS130** | TTGGCGTGGGTGATGCA | 3477187 | + | plu2970 | + | 559 ~~~~~ 798 | |
| **IS131** | AGCAAGATCAACGCCAA | 3487783 | - | plu2985 | - | 385 ~~~~~ 345 | |
| **IS132** | TGCAGTTGTTGCGCCAC | 3490871 | - | plu2988 | - | 509 ~~~~~ 237 | |
| **IS133** | GGCAATACCCGTACCAA | 3492236 | - | plu2990 | - | 304 ~~~~~ 465 | |
| **IS134** | TGCTAATGTCGTGCCAG | 3576897 | - | plu3068 | + | 956 ~~~~~ 482 | |
| **IS135** | AGCAAGATCCCCGCCAA | 3590351 | - | plu3078 | - | 730 ~~~~~ 774 | |
| **IS136** | TGCAATAACAGCGCCAA | 3593064 | - | plu3080 | - | 58 ~~~~~ 228 | |
| **IS137** | CTGGCTCGGTTGTTGCA | 3669557 | + | plu3127 | + | 1563 ~~~~~ 580 | Within BGC |
| **IS138** | GGCAAGTAGCGGGCCAA | 3683221 | - | plu3130 | - | 4693 ~~~~~ 5241 | Within BGC |
| **IS139** | CCGGCATAGCTTCTGCT | 3709258 | + | plu3145 | + | 658 ~~~~~ 327 | Within BGC |

| | | | | | | | |
|---|---|---|---|---|---|---|---|
| **IS140** | TGCAAAAGGCACACCGG | 3717787 | - | plu3158 | - | 264 ~~~~~ 302 | |
| **IS141** | AGCATTTTCTACGCCAA | 3757680 | - | plu3193 | - | 660 ~~~~~ 1015 | |
| **IS142** | AGCAGTGACCGTGCCAT | 3767128 | - | plu3201 | - | 504 ~~~~~ 800 | |
| **IS143** | TTGGCTCGTGATTTGCA | 3774012 | + | plu3206 | + | 114 ~~~~~ 661 | |
| **IS144** | TGCATCAGCGGTGCCAC | 3855435 | - | plu3254 | - | 167 ~~~~~ 352 | |
| **IS145** | TTGGCACGGATGTTGAT | 3856128 | + | plu3255 | - | 319 ~~~~~ 1392 | Within BGC |
| **IS146** | GGCAAATAGCGAGCCAA | 3873015 | - | plu3263 | - | 7923 ~~~~~ 7759 | Within BGC |
| **IS147** | TGCAGATTCCATGCCAA | 3930870 | - | plu3314 | + | 270 ~~~~~ 14 | |
| **IS148** | TGCATAACCTGAGCCAA | 4017637 | - | plu3392 | - | 2005 ~~~~~ 112 | |
| **IS149** | TGCATTAACCATGCCGG | 4032998 | - | plu3413 | - | 120 ~~~~~ 313 | |
| **IS150** | ATGGCCTGATTAATGCA | 4064185 | + | plu3457 | + | 318 ~~~~~ 1464 | |
| **IS151** | TTGGTATCGGTTTTGCT | 4147924 | + | plu3534 | - | 2647 ~~~~~ 7149 | Within BGC |
| **IS152** | AGCAAGTAATACGCCAA | 4156839 | - | plu3535 | - | 1750 ~~~~~ 8091 | Within BGC |
| **IS153** | CTGGCGTAATCCCTGCA | 4176598 | + | plu3544 | - | 185 ~~~~~ 770 | Within BGC |
| **IS154** | AGCAAAATTGGCACCAG | 4196077 | - | plu3561 | - | 502 ~~~~~ 1027 | |
| **IS155** | TGCAATTCTCCTGCCAG | 4214845 | - | plu3575 | - | 343 ~~~~~ 232 | |
| **IS156** | CGCAACATTCACGCCAC | 4275433 | - | plu3623 | - | 2508 ~~~~~ 150 | |
| **IS157** | AGCAATTTCAGTGCCCG | 4334885 | - | plu3680 | - | 142 ~~~~~ 777 | |
| **IS158** | TGCAAACTCAGCGCCAC | 4386145 | - | plu3720 | - | 483 ~~~~~ 193 | |
| **IS159** | GTGGCGTGAGCTTTGCT | 4390158 | + | plu3724 | - | 758 ~~~~~ 763 | |
| **IS160** | CCGGCGTGATTTTTGCG | 4399390 | + | plu3732 | - | 44 ~~~~~ 1406 | |
| **IS161** | ATGGCGCTGAATTTGCA | 4404166 | + | plu3735 | + | 188 ~~~~~ 471 | |
| **IS162** | AGCAAGAACTACGCCGT | 4446423 | - | plu3779 | + | 475 ~~~~~ 777 | |
| **IS163** | GGCATTCTCCGTGCCAG | 4474949 | - | plu3808 | + | 558 ~~~~~ 523 | |
| **IS164** | GGCATGACTTGCGCCAG | 4491663 | - | plu3824 | + | 560 ~~~~~ 2170 | |
| **IS165** | TTGGCAGTATTATTGCT | 4557169 | + | plu3880 | + | 279 ~~~~~ 1095 | |
| **IS166** | AGCAAGAATCGCACCAT | 4575161 | - | plu3902 | - | 1620 ~~~~~ 225 | |
| **IS167** | TGCAGATACTGGGCCAG | 4639331 | - | plu3952 | - | 268 ~~~~~ 568 | |
| **IS168** | TTGGTATCGATTATGCA | 4664624 | + | plu3978 | + | 836 ~~~~~ 903 | |
| **IS169** | GTGGAGCAGATGTTGCA | 4678188 | + | plu3990 | + | 1956 ~~~~~ 60 | |

| | | | | | | | |
|---|---|---|---|---|---|---|---|
| **IS170** | TGCAATGTCTGCGCCAT | 4719682 | - | plu4027 | + | 1416 ~~~~~ 138 | |
| **IS171** | TGCAATATCCGCACCCG | 4720705 | - | plu4028 | - | 444 ~~~~~ 805 | |
| **IS172** | AGCAGGATCTGCGCCAG | 4792770 | - | plu4104 | - | 411 ~~~~~ 863 | |
| **IS173** | TTGGCATTGGATTTGCC | 4827506 | + | plu4138 | + | 77 ~~~~~ 1221 | |
| **IS174** | CTGGCGCAAATAATGCC | 4869827 | + | plu4168 | - | 465 ~~~~~ 4225 | |
| **IS175** | AGCAAAGTTTGTACCAT | 4874599 | - | plu4169 | - | 448 ~~~~~ 2436 | |
| **IS176** | TGCATCAGTAATGCCAG | 4911764 | - | plu4198 | - | 64 ~~~~~ 1311 | Within BGC |
| **IS177** | TGCATCAACGGTGCCAC | 4926091 | - | plu4211 | - | 136 ~~~~~ 339 | |
| **IS178** | CGCAAAGACTGCGCCAG | 4929727 | - | plu4214 | - | 2536 ~~~~~ 807 | |
| **IS179** | TGCAGGCTCTGCGCCAG | 4929820 | - | plu4214 | - | 2629 ~~~~~ 714 | |
| **IS180** | GGCAATCACTGCGCCAA | 4948659 | - | plu4227 | - | 993 ~~~~~ 523 | |
| **IS181** | TTGGCGCGGGTATTGCA | 4955380 | + | plu4232 | + | 40 ~~~~~ 1404 | |
| **IS182** | TTGGCGCAACAGTTGCT | 4991297 | + | plu4266 | + | 49 ~~~~~ 1062 | |
| **IS183** | CTGGAACGACAAATGCA | 4995440 | + | plu4269 | + | 774 ~~~~~ 672 | |
| **IS184** | AGCAACACATATGCCAA | 5034376 | - | plu4308 | + | 34 ~~~~~ 780 | |
| **IS185** | AGCATTAACTATGCCGA | 5044134 | - | plu4319 | + | 180 ~~~~~ 1372 | |
| **IS186** | TTGGCACGACCATTGCT | 5045137 | + | plu4319 | + | 1183 ~~~~~ 369 | |
| **IS187** | TGCATAAACCCTGCCAC | 5148045 | - | plu4401 | - | 274 ~~~~~ 1059 | |
| **IS188** | GTGGCACTAGCTTTGCT | 5213809 | + | plu4458 | + | 1496 ~~~~~ 723 | |
| **IS189** | TGCAGAAATTACGCCAA | 5267257 | - | plu4508 | + | 2053 ~~~~~ 696 | |
| **IS190** | TGCAAATAGATTGCCAG | 5278480 | - | plu4520 | + | 960 ~~~~~ 66 | |
| **IS191** | AGCAACAAACGCGCCAA | 5284826 | - | plu4525 | - | 210 ~~~~~ 1914 | |
| **IS192** | ATGGCCCGAAGCTTGCA | 5314154 | + | plu4553 | - | 1095 ~~~~~ 2662 | |
| **IS193** | AGCAACTTGTACGCCAC | 5314349 | - | plu4553 | - | 1290 ~~~~~ 2467 | |
| **IS194** | TGCAACATCTGTTCCAT | 5339438 | - | plu4575 | - | 607 ~~~~~ 1831 | |
| **IS195** | TGCATAGACGGTGCCAG | 5345756 | - | plu4580 | - | 35 ~~~~~ 1243 | |
| **IS196** | CAGGAACGGATCTTGCA | 5347371 | + | plu4581 | - | 327 ~~~~~ 32 | |
| **IS197** | GGCAAAATTTACGCCGA | 5393448 | - | plu4626 | + | 672 ~~~~~ 124 | Within BGC |
| **IS198** | AGCATTTTCTATGCCAA | 5428795 | - | plu4656 | - | 474 ~~~~~ 100 | |
| **IS199** | AGCAAGAACAGTGCCAA | 5434703 | - | plu4662 | - | 100 ~~~~~ 969 | |

| | | | | | | |
|---|---|---|---|---|---|---|
| **IS200** | AGCAAAACAGATGCCAA | 5435503 | - | plu4662 | - | 900 ~~~~~ 169 |
| **IS201** | GTGGCACAAACTGTGCA | 5453601 | + | plu4676 | + | 1237 ~~~~~ 27 |
| **IS202** | AGCAATTGCCTTGCCAA | 5490617 | - | plu4710 | - | 117 ~~~~~ 220 |
| **IS203** | TGCAAAAGTTACGCCAC | 5497303 | - | plu4725 | - | 348 ~~~~~ 252 |
| **IS204** | AGCAATAACAATGCCAA | 5503995 | - | plu4732 | - | 486 ~~~~~ 457 |
| **IS205** | GGCAAAACACACGCCAG | 5558998 | - | plu4777 | - | 516 ~~~~~ 156 |
| **IS206** | TGCATTACCTGCGCCAA | 5572888 | - | plu4799 | + | 164 ~~~~~ 901 |
| **IS207** | TAGGCATCGATATTGCA | 5602417 | + | plu4826 | + | 33 ~~~~~ 147 |
| **IS208** | AGCATTAGCTATGCCAT | 5622307 | - | plu4848 | + | 158 ~~~~~ 875 |
| **IS209** | CTGGCGCATATGTTGCA | 5642081 | + | plu4868 | + | 57 ~~~~~ 564 |
| **IS210** | CTGGTACTGATATTGCT | 5658182 | + | plu4887 | + | 241 ~~~~~ 2202 |
| **IS211** | TTGTCACGGATTTTGCA | 5661493 | + | plu4889 | + | 156 ~~~~~ 469 |

[a]Biosynthetic gene cluster boundaries as determined by antiSMASH.

**Table S5.** Proteins encoded by *Photorhabdus laumondii* TTO1 that have significant sequence similarity to the AAA+ ATPase core of *S. meliloti* DctD.

| **Name** | | **(Bits)** | **Value** | **Identity** |
|---|---|---|---|---|
| WP_011147503.1 | Transcriptional regulator; glrR | 194 | 1.00E-59 | 99/192 |
| WP_011144634.1 | Nitrogen regulation protein NR(I); glnG | 167 | 2.00E-49 | 87/189 |
| WP_011146808.1 | Psp operon transcriptional activator; PspF | 162 | 1.00E-48 | 87/200 |
| WP_011145558.1 | Signal transduction histidine kinase. PAS-containing. | 166 | 2.00E-48 | 84/193 |
| WP_011146802.1 | Transcriptional regulatoryprotein; TyrR | 162 | 2.00E-47 | 83/200 |
| WP_011147723.1 | Propionate catabolism operon regulatory protein; PrpR | 149 | 1.00E-42 | 83/187 |
